# Scalable image-based visualization and alignment of spatial transcriptomics datasets

**DOI:** 10.1101/2021.12.07.471629

**Authors:** Stephan Preibisch, Michael Innerberger, Daniel León-Periñán, Nikos Karaiskos, Nikolaus Rajewsky

## Abstract

We present STIM, an imaging-based computational framework focused on visualizing and aligning high-throughput spatial sequencing datasets. STIM is built on the powerful, scalable ImgLib2 and BigDataViewer (BDV) image data frameworks and thus enables novel development or transfer of existing computer vision techniques to the sequencing domain characterized by datasets with irregular measurement-spacing and arbitrary spatial resolution, such as spatial transcriptomics data generated by multiplexed targeted hybridization or spatial sequencing technologies. We illustrate STIM’s capabilities by representing, interactively visualizing, 3D rendering, automatically registering and segmenting publicly available spatial sequencing data from 13 serial sections of mouse brain tissue, and from 19 sections of a human metastatic lymph node.

## Introduction

Several recent technological breakthroughs have triggered the rapid development of numerous high-throughput spatial transcriptomics methods over the last few years. These fluorescent RNA hybridization-based^1–5^ or array-based RNA capture, barcoding, and subsequent sequencing^6–13^ techniques provide (typically at single-cell or subcellular resolution) molecular readouts within the native spatial context of a tissue, which is critical for understanding cellular interactions in healthy and diseased states^14^. Therefore, the production of spatial transcriptomics datasets, already abundant and continuously expanding, is crucial for both basic life science research and clinical/medical applications. Handling and analysis of these datasets pose several complex challenges: (a) size – spatial sequencing generates several gigabytes of data for a single tissue section and we anticipate that to increase; (b) heterogeneity – datasets greatly differ in the number of genes and transcripts captured, their spatial resolution, and tissue architecture; (c) spatial transcriptomics data are in contrast to image data usually irregularly spaced; (d) three-dimensional (3D) integration – data from tissue sections need to be integrated into a 3D molecular map; (e) access and analyses – the need to easily share and interactively interrogate spatial transcriptomics data, and (f) flexibility and long-term availability – the need for open source, community-based approaches.

Several methods have been developed to visualize, process, and align spatial transcriptomics data^15–31^ – each however having their own drawbacks. Here, we show that powerful methods from the computer vision field that have been developed by a large scientific community for decades, can alternatively be adapted to meet the challenges that spatial transcriptomics methods face. Specifically, we present the “Spatial Transcriptomics Imaging Framework” (STIM), a computational, scalable and extendable toolkit based on ImgLib2^32^ that allows efficient handling, processing (including integration of 2D data into 3D molecular maps), visualization, alignment and analysis of high-throughput spatial -omics datasets. We demonstrate the power of our approach by applying STIM to two distinct spatial sequencing datasets, integrating adjacent slices into 3D molecular maps: (i) adult mouse brain, and (ii) a challenging dataset of 19 sections from a human metastatic lymph node. We additionally show how to use STIM to visualize the data and perform a simple, machine learning-based segmentation task.

## Results

ImgLib2^32^ defines an image as a function *f* that maps coordinates *C* in n-dimensional space *R*^*n*^ to a value *T*

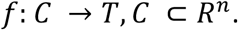

This definition illustrates that ImgLib2 natively supports both regularly and irregularly spaced datasets (**Sup. Fig. 1**). Furthermore, its generic, interface-driven design imposes no constraints on dataset size (biggest currently available implementation supports 4,096 petabyte), dimensionality or data type, which is highlighted by the fact that many of the largest biological image datasets ever acquired^33,34^ were reconstructed using ImgLib2, the ImgLib2-backed BDV^35,36^ and the N5 (zarr-compatible) file format^37^. STIM builds on these frameworks to provide fast, random and distributed read&write access, interactive visualization and efficient processing of spatial transcriptomics data.

First, STIM directly supports AnnData or re-saving of input datasets from the standardized text- or comma-separated formats into an N5 container (**Fig 1a,b**) while optionally log-normalizing the data. Coordinate- and gene expression data can be loaded fast and memory-efficiently in blocks using the Imglib2-cache framework, which can be accessed as values or as rendered images (**Fig. 1b,c**). ImgLib2 provides nearest-neighbor and linear interpolation for mapping irregularly-spaced samples onto pixel grids necessary for visualization. For a more realistic rendering at arbitrary resolutions we implemented a rendering method based on Gaussian distributions (**Fig. 1, Methods, Sup. Methods**). Spatial image filtering, also referred to as digital filtering, is an established, powerful technique to enhance certain aspects of a signal that is represented as discrete samples (e.g. an image) using mathematical operations^38^. While such filtering is mathematically directly applicable to irregularly-spaced data, it is not widely available as efficient implementations require fast k-nearest neighbor search. To ease this barrier, we added a generic framework based on ImgLib2 using kd-trees for applying filters (e.g. Mean, Median, Gaussian) to irregularly-spaced data that can easily be extended (**Fig. 1c**). All operations are implemented virtually allowing interactive access to and rendering of the data using BDV.

**Figure 1:**
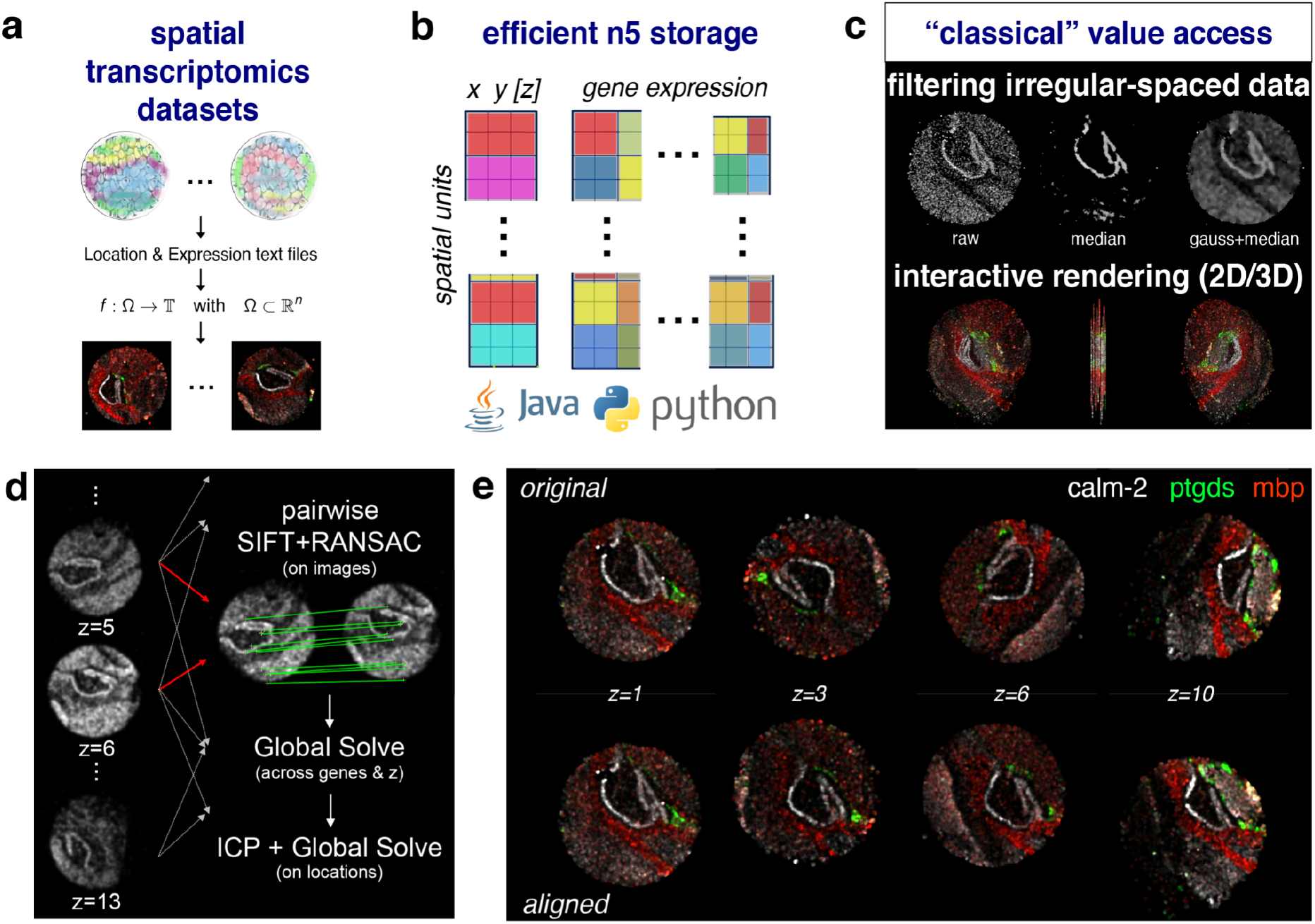
STIM applies state-of-the-art imaging techniques to store, visualize, and analyze massive amounts of spatial transcriptomics datasets. **a**, Spatial transcriptomics datasets can be represented as images with the number of genes per spatial unit corresponding to a number of different colors. **b**, Expression and spatial data is stored in N5 containers for efficiency and scalability and is accessible within the provided Java and Python frameworks. **c**, STIM provides classical value access operations such as filtering irregular-spaced data, resulting in smoothed gene expression, or interactive rendering in 2D and 3D. **d**, Schematic of how STIM aligns consecutive sections of spatial transcriptomics datasets. **e**, Visualization of the spatial gene expression of three genes in four different sections of a published dataset. Top (bottom) row: spatial gene expression profiles in the original (aligned) puck orientations.

We demonstrate STIM’s capabilities as follows. First, STIM can be used to filter the data and smoothen it, for instance by applying a Median filter or others (**Fig. 1c**). Second, tried-and-tested image registration techniques can be used to align datasets stemming from consecutive sections of the same tissue as described below (**Fig. 1d,e**). Of note, we also developed a user-friendly BDV-based GUI for aligning tissue sections interactively using SIFT and ICP, or, optionally, manually using standard BigDataViewer transformation controls **(Sup. Fig. 7**). Third, STIM offers an interactive visualization and exploration of the data through BDV in 2D and 3D, including the visualization of metadata -- such as cell type annotation -- together with gene expression in every spatial unit (**Sup. Fig. 2, Sup. Movie 2**). Additionally, we show that existing machine learning segmentation can be straightforward applied to spatial transcriptomics data and we highlight the applicability of existing 3D rendering methods (**Fig. 2/Sup. Movie, Sup. Methods**).

**Figure 2 (Movie, submitted as Sup. Movie 1):**
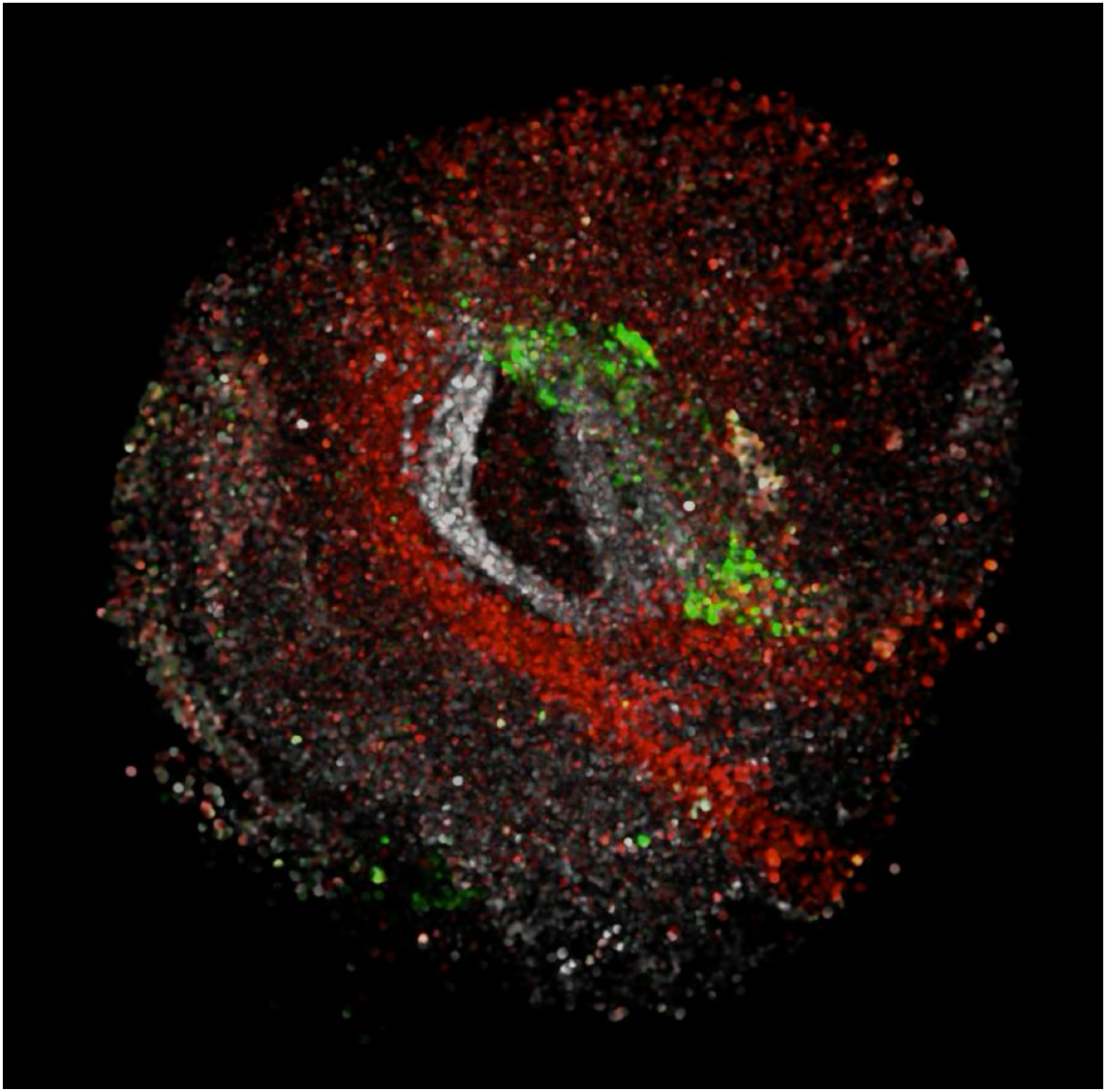
3D rendering of the aligned SlideSeq dataset. The movie shows gene Calm2 is shown in white, gene Ptdgs in green, and gene Mbp in red and highlights the 3D nature of the dataset after alignment when represented as an image.

### Alignment

To illustrate the potential of applying computer vision techniques to sequencing datasets, we aligned a series of consecutive brain cerebellum sections using STIM that were published in ^7^ (**Fig. 2 / Sup. Video 1**). Each of these 2D sections contains between 12,000 and 33,000 cells and a median of ∼50 quantified molecules at near-cellular resolution. To align the 13 sections in three dimensions we adapt an alignment strategy originally developed for the registration of large electron and light microscopy datasets^36,39,40^. We first apply Scale Invariant Feature Transform (SIFT)^41^ in combination with Robust Sample Consensus (RANSAC)^42^ on rendered image pairs of sections (+-2 in the z-direction, **Fig. 1d**). For each pair, we identify a set of corresponding points on a rigid 2D transformation across an automatically selected set of genes that show high entropy in both sections. To correctly identify as many corresponding points as possible we run SIFT independently for each gene with a low threshold for the minimal number of required points. RANSAC is then applied to all points again to identify those that agree on a common transformation across genes. Next, we globally minimize the distance between all corresponding image points across all sections, yielding a single 2D transformation for each section (**Sup. Methods**). In an optional refinement step, we use the Iterative Closest Point (ICP) algorithm^43^ on locations of sequenced spots rather than rendered images, where neighboring points within a predefined radius showing most similar expression values are assigned to be correspondences. Finally, using all ICP correspondences, we globally solve again and identify a regularized affine transformation model for each section that is stored in the N5 container. The resulting tissue dataset can be rendered in 3D using STIM (**Fig. 2 / Sup. Video 1**) and the STIM-explorer can be used to highlight the spatial expression of interactively selected genes on the whole tissue (**Sup. Video 2**).

Due to the robustness of SIFT, our alignment pipeline can readily be employed across different technologies. Importantly, STIM was vital in constructing a 3D molecular map of a recently published human metastatic lymph node^13^. This challenging dataset consists of 19 non-consecutive sections and a total of >1.5 million cells. The pairwise and global alignment performed by STIM enabled the generation of the 3D virtual tissue block, and the derivation of 3D-specific insights from the data^13^. In another application, we used STIM to align six sections of human lung cancer tissue, each containing ∼50,000 cells and being 30 μm apart^44^. Among other insights, the 3D molecular map in that case enabled the more precise identification of immune niches^44^.

We further used STIM to align serial adult mouse brain sections produced with the Visium platform (**Sup. Fig. 3**). More generally, we anticipate STIM to process and stitch together consecutive tissue sections from the same tissue, regardless of the underlying spatial sequencing method. This enhances the information flow between the different sections and naturally enriches the molecular readouts.

The alignment pipeline we developed is based on linear transformations, typically affine transformation models regularized with rigid models. This allows the use of robust model estimation using RANSAC, which is also able to realize if no proper alignment could be achieved. If the alignment quality needs to be further improved, existing non-rigid registration algorithms such as bUnwarpJ^39,45^ can be employed. However, it is important to realize that while deformations introduced by non-rigid transformations typically do improve the alignment quality (**Sup. Fig. 4**), it is likely to deform the sample in an unnatural way. A simple example to visualize the problem is a cone in 3D, which would be represented as circles with increasing radius along the sections (in z). Non-rigid alignment would effectively transform these circles so they show the same radius as it maximizes similarity, thus transforming the cone into a pipe. Therefore, meaningful and well-designed regularization is a necessity for employing non-rigid alignment, which has been well-studied in image analysis^39,45^ but is, to the best of our knowledge, currently in its infancy for ST data.

### Interoperability and accessibility

STIM is open-source and leverages the large Java community built around ImgLib2. To enhance interoperability and enable the use of STIM by users who employ Python interfaces, we have added support for the popular AnnData format^46^. With this support, it is possible to seamlessly access the data between the AnnData and the n5 formats, transferring the underlying sample metadata, and facilitating downstream analyses. Moreover, STIM can be installed on Linux, MacOS, and Windows through the popular Conda packaging environment.

### Benchmarking

Recently, several software packages for the alignment of spatial sequencing data have been developed specifically within the field of spatial transcriptomics (**Sup. Table 1**). Probabilistic Alignment of Spatial Transcriptomics Experiments (PASTE)^47^ first solves an optimal transport problem to derive a probabilistic assignment of points for pairs of consecutive slices. Based on these, a rigid transformation model is sequentially estimated for each slice. Importantly, no optimization across slices is performed, which might prove problematic once datasets increase (similar to the image stitching problem), and *partial* alignment (i.e., sections only partially overlap in 2D) is not supported. More recently, PASTE2 ^31^ introduced support for partial alignment, but the optimal transport framework is not scalable (in time or memory usage) to datasets with millions of cells. Furthermore, the computational complexity depends cubically on the number of sequenced locations. Andersson et al.^23^ relies on manual landmarks for alignment and also supports only rigid transformation models. Jones et al. ^24^ and Qiu et al.^25^ require an approximate initial alignment, using for example PASTE or STIM, and apply Gaussian Process Spatial Alignment to extract warp functions for non-rigid alignment of consecutive slices, also at cubic complexity with respect to the number of sequenced locations. Clifton et al.^26^ require initialization via manual selection of corresponding points, and do not offer global optimization across slices. Other methods rely on alignment of high-level features such as cell types or spatial regions, thus requiring extensive analysis of individual sections prior to alignment^27–29^. Our proposed alternative approaches have been tried-and-tested in image analysis for decades, work reliably and fast on multi-terabyte images while their complexity depends on the size of the rendered images, which can even be small for ST data. The complexity for transferring sequenced locations into images is equivalent to that of a kdTree lookup, which is ***O(log n)***. The use of RANSAC for pairwise matching, global optimization with outlier removal across all pairwise results, as well as implementation in scalable frameworks ImgLib2 and BigDataViewer ensures that the identified alignment can be trusted and that the approach will scale to significantly larger datasets in the future.

## Discussion

STIM enables efficient, distributed access, processing and visualization of large-scale spatial transcriptomics datasets. Irregularly-spaced data can be spatially filtered and accessed directly as values or rendered as images. STIM thereby acts as a bridge between the fields of computer vision and genomics, which we highlight by developing an automatic workflow for the alignment of sliced spatial transcriptomics datasets. Another application of STIM is to perform object segmentation on subcellular, high-resolution spatial transcriptomics datasets using existing image-based machine learning solutions such as Random Forests (**Sup. Fig. 5**)^48^ or for larger future datasets StarDist^49^ or CellPose^50^. We provide STIM as an extensible open source framework available on GitHub with interfaces in Java, Python and on the command line. We believe that these properties of STIM have the potential to enable the community to further unite the worlds of image analysis and genomics.

## Methods

### Related software

Spacemake ^30^ is used for processing and basic visualization only; VT3D^51^, Spateo Viewer^25^, and Vitessce^15^ are designed for visualization purposes only; Seurat^16^ and squidpy^17^ offer both analyzing and viewing, but 3D visualization and sections alignment are unavailable; ST Viewer^18^ is tailored to datasets generated with Spatial Transcriptomics^6^; histoCAT^19^ and Giotto Viewer^20^ do not offer sections alignment. Semla^52^ (formerly STUtility^21^) offers a basic alignment, but requires manual registration for high-quality results.

### N5 storage and normalization

STIM ingests spatial sequencing data that is stored in the AnnData standard, or as (compressed) text files containing locations and barcodes of sequenced spots, expression levels, and optionally cell type predictions per dataset. The current framework supports 2D and 3D coordinates, and can be readily extended. Initially, STIM re-saves or links the set of AnnData or text-file based datasets into a common N5 container that can be accessed as one common project, e.g. the image registration pipeline can be applied to an entire or parts of a project. By default, locations and expressions are stored with double precision and Gzip compression using a block length of 16,384 for locations and a block size of 512×512 for expression values (Fig. 1b). Genes and the barcode list of each dataset together with their transformations are currently stored as metadata in the N5 container. The N5 data can be accessed in STIM/stimwrap or directly through Java and python N5 packages.

Optionally, the data can be normalized upon re-saving to N5 or at a later time. We have adopted here the standard library size normalization in log-space commonly used for scRNA-seq datasets. More specifically, if *d*_*ij*_ represents the raw count for gene *i* in spatial unit *j*, we normalize values as

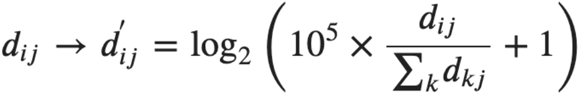

where the dummy index *k* is used for summing over all genes within a spatial unit.

### Rendering of irregularly-spaced data

Image data is typically stored in n-dimensional, integer-based cartesian coordinate systems that are natively supported by most camera chips and display devices. In contrast, spatial sequencing data consist of measurements at arbitrary, floating-point precise locations, which stem from accurately localizing sequenced locations. To render such data, they need to be mapped to an integer-based cartesian coordinate system supported by standard display devices (e.g. to a 2048×1536 pixel grid), a problem that also occurs in other dataset types such as localization-based superresolution microscopy^53^ and other disciplines such as astronomy^54^. Straight-forward, fast mapping can be achieved through nearest-neighbor interpolation using kd-trees. Resulting images are effectively Voronoi-tessellations with sharp, unnatural boundaries and artificial appearance where point densities are low towards the edges (**Sup. Fig. 6A,B**). Distance-weighted interpolation creates more natural-looking images, which, however, still contain unnatural edges as either a number of points or maximal distance for interpolation needs to be defined (**Sup. Fig. 6C**). Large values are able to create reasonable representations in areas of high point densities, but can still produce artificial structures, especially towards the edges of the dataset (**Sup. Fig 6D**). To overcome these issues we represent each location as a Gaussian distribution, and each pixel is rendered as the sum of all overlapping distributions, normalized by their respective weights. In order not to create hard boundaries in areas with few locations, normalization is only performed if the sum of weights is bigger than one. This mapping is fast and produces representations of the data resembling naturally-looking images that can therefore easily be processed with computer vision tools and are additionally visually pleasing (**Fig. 1, Sup. Video 2, Sup. Fig. 6E**).

### Filtering of irregularly-spaced data

Within STIM, we implemented a framework for spatial filtering of irregularly-spaced datasets based on kd-trees. We added mean filtering, median filtering, Gaussian filtering, as well as practical filters to hide single, isolated locations and to visualize the density of locations. Adding new filters is straight-forward, and typically requires the implementation `filter.RadiusSearchFilter` class, which already provides the kd-tree search and the location to be filtered.

Such basic filtering operations can for example help to smoothen noisy spatial sequencing data, to emphasize larger structures, or to identify edges (**Fig. 1c, Sup. Video 2**).

### Pairwise SIFT registration

In order to robustly identify corresponding points between pairs of two-dimensional serial sections of spatial sequencing datasets we first employ the Scale Invariant Feature Transform (SIFT) on images of renderings of individual genes.

First, we identify a set of genes (by default 100) that are expressed in both serial sections and show the highest combined standard deviations of their expression values, thus automatically selecting genes that are likely to show patterns that are helpful to perform an alignment. The user can additionally add genes that are known to create well-structured expression renderings. Second, we compute SIFT on all pairs of genes individually, using a low minimal number of corresponding points (inliers, by default 5) on a rigid model. Finally, we perform another RANSAC consensus across the points of all genes, requiring by default at least 30 inliers. This combination of parameters was very robust in our tests and worked out of the box for SlideSeq, Visium, and Open-ST datasets.

### Global Optimization

In order to align more than two serial sections we first compute pairwise SIFT registrations between close-by sections (by default +-2). We then solve an optimization problem by finding a set of transformations *T*_*V*_ that minimize the distance between all corresponding points *C*_*A,B*_(*g*) of all serial sections *V* across all genes *G* by identifying ^36^.

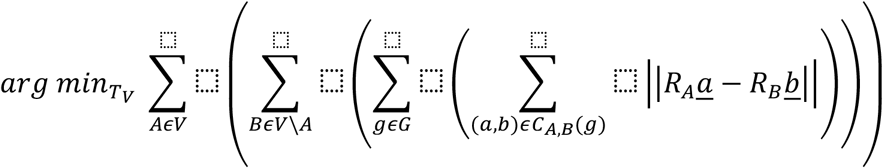

Finally, wrong pairs of correspondences *C*_*A,B*_(*g*) can be identified and removed by iteratively analyzing inconsistencies between pairwise results and the current state of the global optimization as defined by the current set of transformations *T*_*V*_^40^. By default we employ rigidly regularized (*α* = 0.1) affine transformation models *T*_*V*_ for each serial section in the optimization process. All transformations are stored within the N5 metadata, thus all visualization tools of STIM will directly use these transformations.

### ICP refinement

After global optimization based on the corresponding interest points identified by SIFT we optionally employ Iterative Closest Point (ICP) for refinement of the transformations. In contrast to the SIFT alignment step, ICP is performed on the actual coordinates of the sequenced locations. We first compute pairwise ICP’s between close-by serial sections (+-2 sections) using only the expression values of genes that yielded SIFT correspondences. The basic idea of ICP is to assign nearest neighboring points as corresponding points, update the transformation based on this assignment, and iterate this procedure until convergence or a maximum number of iterations is achieved. Here, we do not simply assign nearest points to each other, but those who show the most similar expression vector in the local vicinity (by default the median distance between all sequenced locations). We optionally support RANSAC filtering on the sets of corresponding points during each ICP iteration in order to identify a consensus update vector across all neighboring points. After all pairwise ICP matchings are performed, we re-solve the global optimization problem using the corresponding points identified in the last iteration of every respective ICP run.

## Data availability

All data analyzed within this work are publicly available. The datasets that were used for the alignment of mouse cerebellum sections were published in Slide-seq^7^. The 10X Visium datasets were downloaded from the 10X Genomics website. The Open-ST data were downloaded from GEO (GSE251926).

## Code availability

STIM and stimwrap are available on GitHub (https://github.com/PreibischLab/STIM and https://github.com/rajewsky-lab/stimwrap)

## Acknowledgements

S.P. was supported by the HFSP grant RGP0021/2018-102, MDC Berlin and HHMI Janelia. M.I. was supported by HHMI Janelia. D.L.-P. was supported by the Helmholtz Einstein International Berlin Research School in Data Science (HEIBRiDS) program of the Helmholtz Association. N.K. was supported by the DFG grants KA 5006/1-1 and RA 838/5-1. N.R. was supported by MDC Berlin and Charite. We thank the HHMI Janelia Open Science Software Initiative (OSSI, https://ossi.janelia.org/) for supporting this project.

## Declaration of interests

This work is part of a larger patent application in which N.K., S.P. and N.R. are among the inventors. The patent application was submitted through the Technology Transfer Office of the Max-Delbrück Center (MDC), with the MDC being the patent applicant.

## Supplementary Notes

### Supplementary Figures

1. Comparison of regularly and irregularly spaced datasets
2. Interactive overlay of cell type onto a SlideSeq dataset
3. Alignment of a 10x Visum dataset
4. Comparison of non-rigid and regularized affine alignment
5. Applying existing machine learning segmentation software to spatial transcriptomics
6. Comparison of different point cloud rendering methods supported by STIM
7. Interactive alignment using SIFT in the STIM BigDataViewer-based GUI

### Supplementary Tables

1. Comparison of 3D registration methods for Spatial Transcriptomics data

### Supplementary Videos

1. 3D rendering of the aligned SlideSeq dataset, gene *Calm2* is shown in white, gene *Ptdgs* in green, and gene *Mbp* in red.
2. Screen recording of the st-explorer for gene *Calm2* of the SlideSeq dataset. Highlighted are interactive exploration, as well as different levels of median filtering on irregularly spaced data, as well as different values for the sigma of the Gaussian point cloud rendering.
3. Exemplary run through the STIM alignment GUI highlighting major features

### Supplementary Notes

1. Workflow for the alignment of the 10x Visium dataset
2. 3D Rendering of the SlideSeq dataset

**Supplementary Figure 1:**
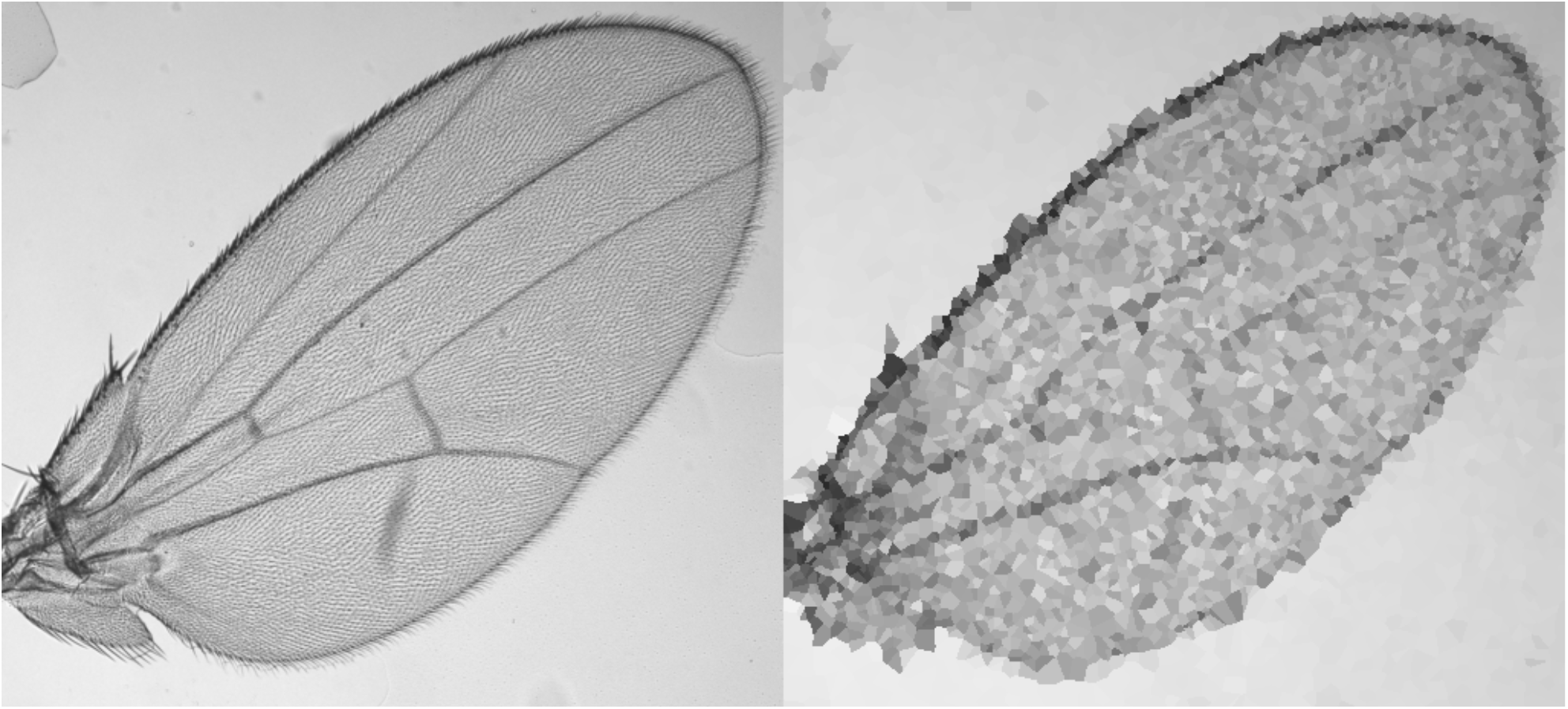
Comparison of regularly and irregularly spaced datasets. The left panel shows a regular image of a Drosophila wing captured by a standard widefield microscope (image courtesy of Prof. Nicolas Gompel). The right panel shows was created by extracting 5000 points at random subpixel locations from the original image and rendering it using the built-in nearest neighbor interpolation for sparse datasets in ImgLib2.

**Supplementary Figure 2:**
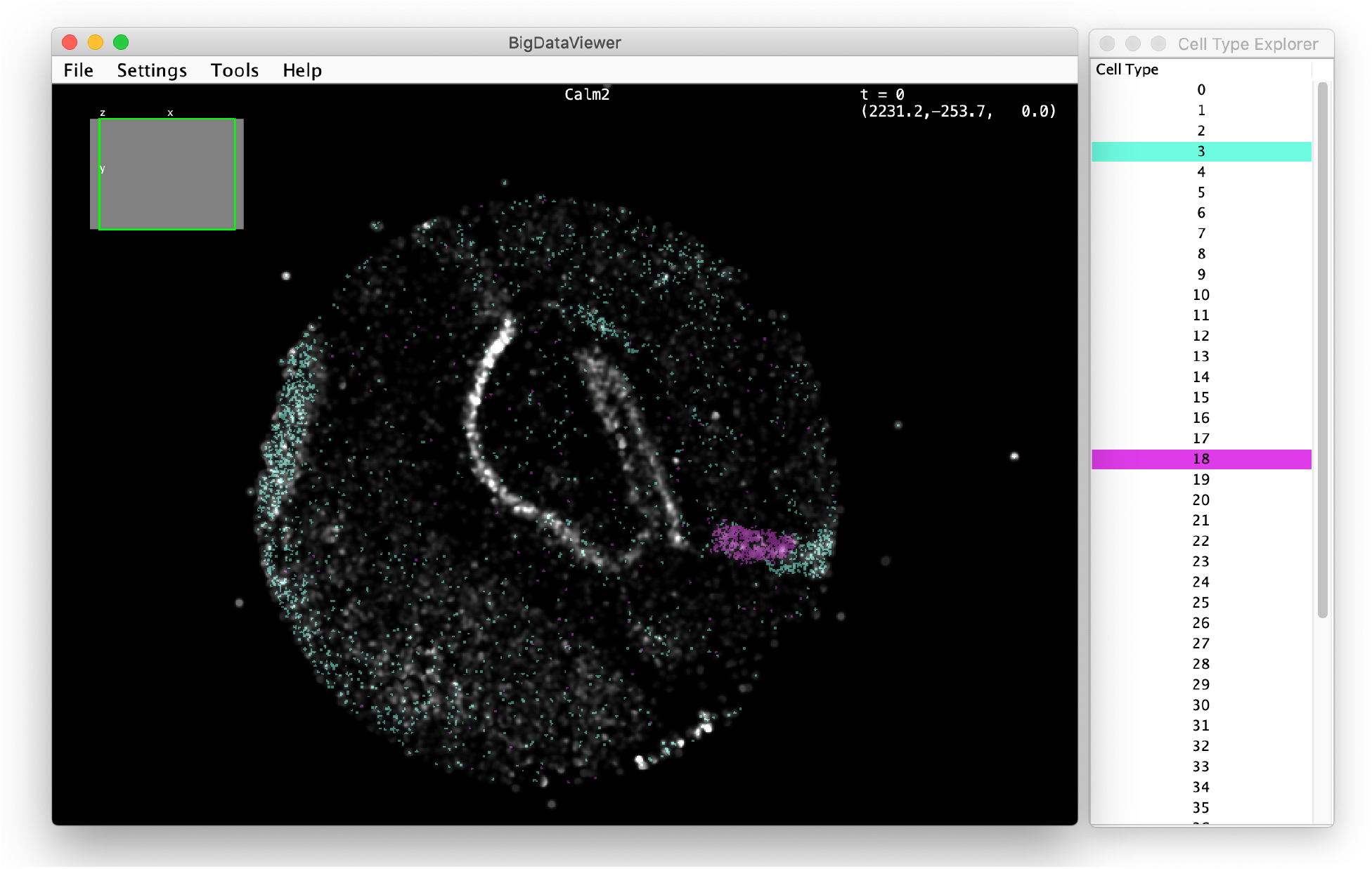
Interactive overlay of cell type onto a SlideSeq dataset. Depicted is a screenshot using `st-bdv-view` that visualizes the gene *Calm2* (more than one gene can be displayed in parallel), together with all predicted cell types. Two of them are interactively highlighted. The screenshot was created calling: `./st-bdv-view -i /home/slide-seq.n5/ -d Puck_180531_22 -g Calm2 -rf 1.0 -a celltype`

**Supplementary Figure 3:**
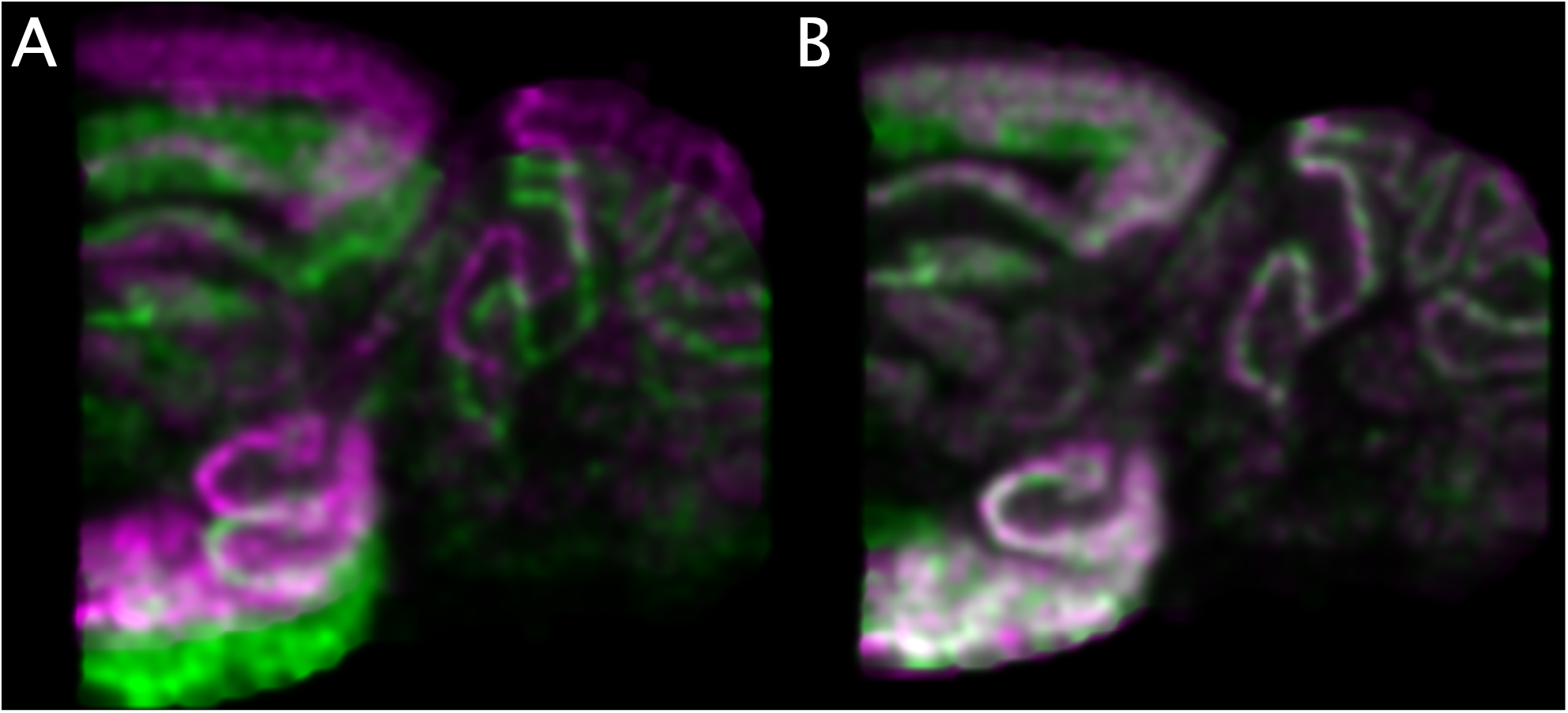
Alignment of a 10x Visium dataset. (**A**) shows the Gaussian point cloud rendering of the gene *Calm2* before alignment, (**B**) shows the same gene after applying an affine transformation. The alignment and visualization were performed on the command line as outlined below. In this case ICP refinement did not improve the alignment quality further and was therefore omitted.

**Supplementary Figure 4:**
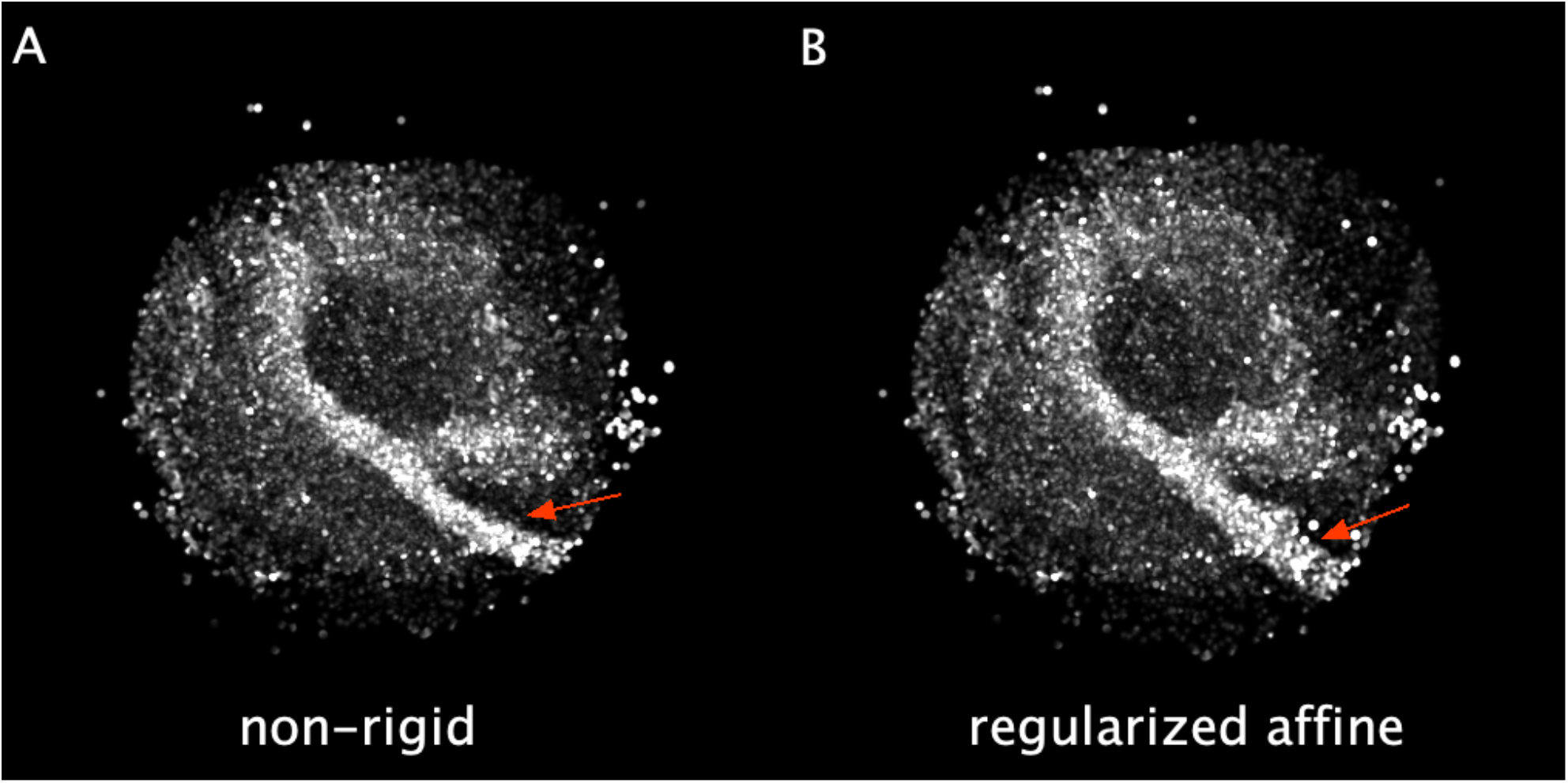
Comparison of non-rigid and regularized affine alignment. (**A,B**) show maximum intensity projections of 8 selected slices of the SlideSeq dataset for the gene *Mbp*. The other slices were omitted due to insufficient data quality that did not allow for an automatic non-rigid alignment. (A)

**Supplementary Figure 5:**
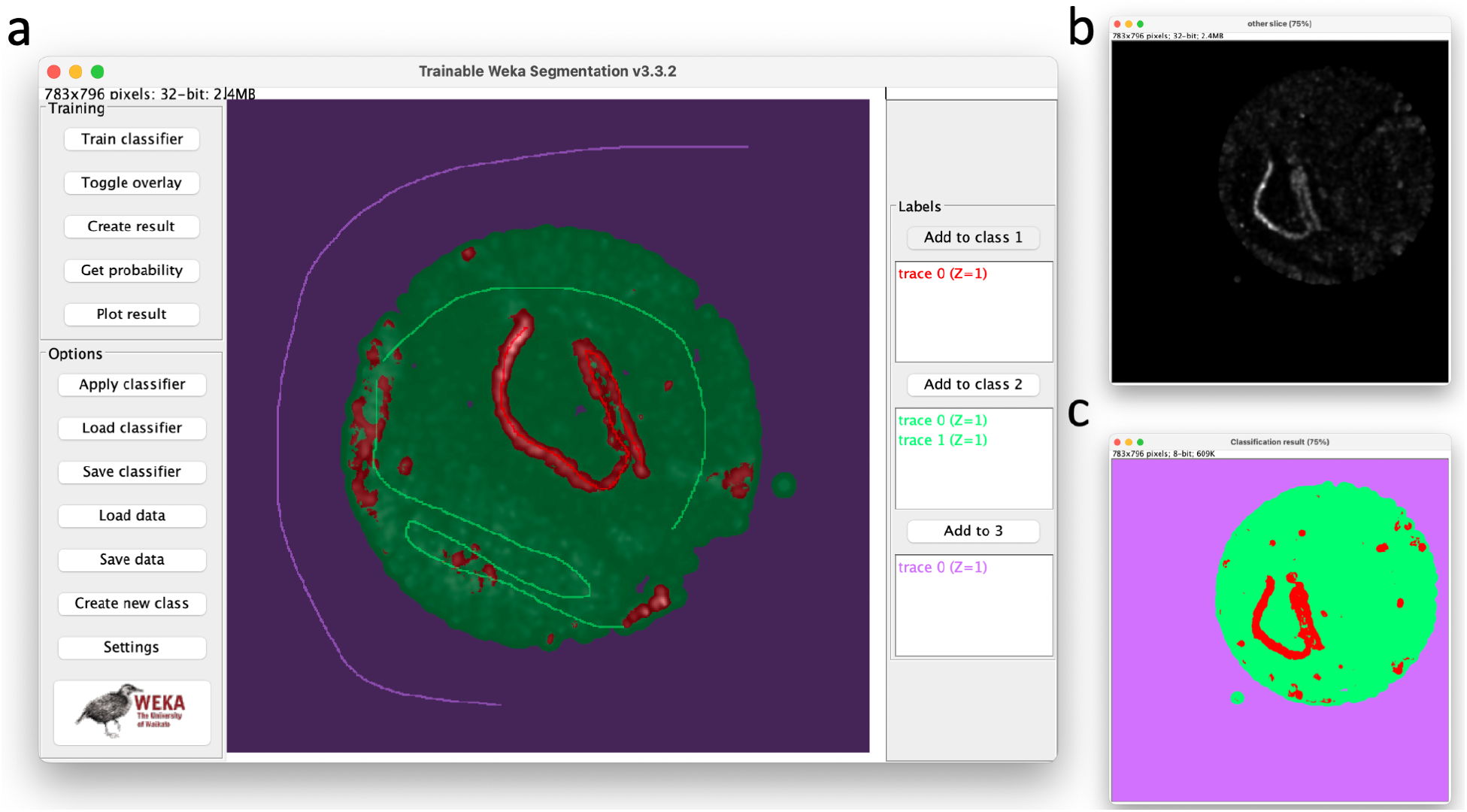
Applying existing machine learning segmentation software to spatial transcriptomics. (**a**) illustrates how the Trainable Weka Segmentation (Random Forest-based method) is used to manually annotate a prominent visible structure in a Slide-Seq dataset using the *Calm2* gene. Note that the annotation effort is limited to the clearly visible lines belonging to each class. (**b**) shows another slice of the dataset to whom the trained classifier was applied (**c**).

**Supplementary Figure 6:**
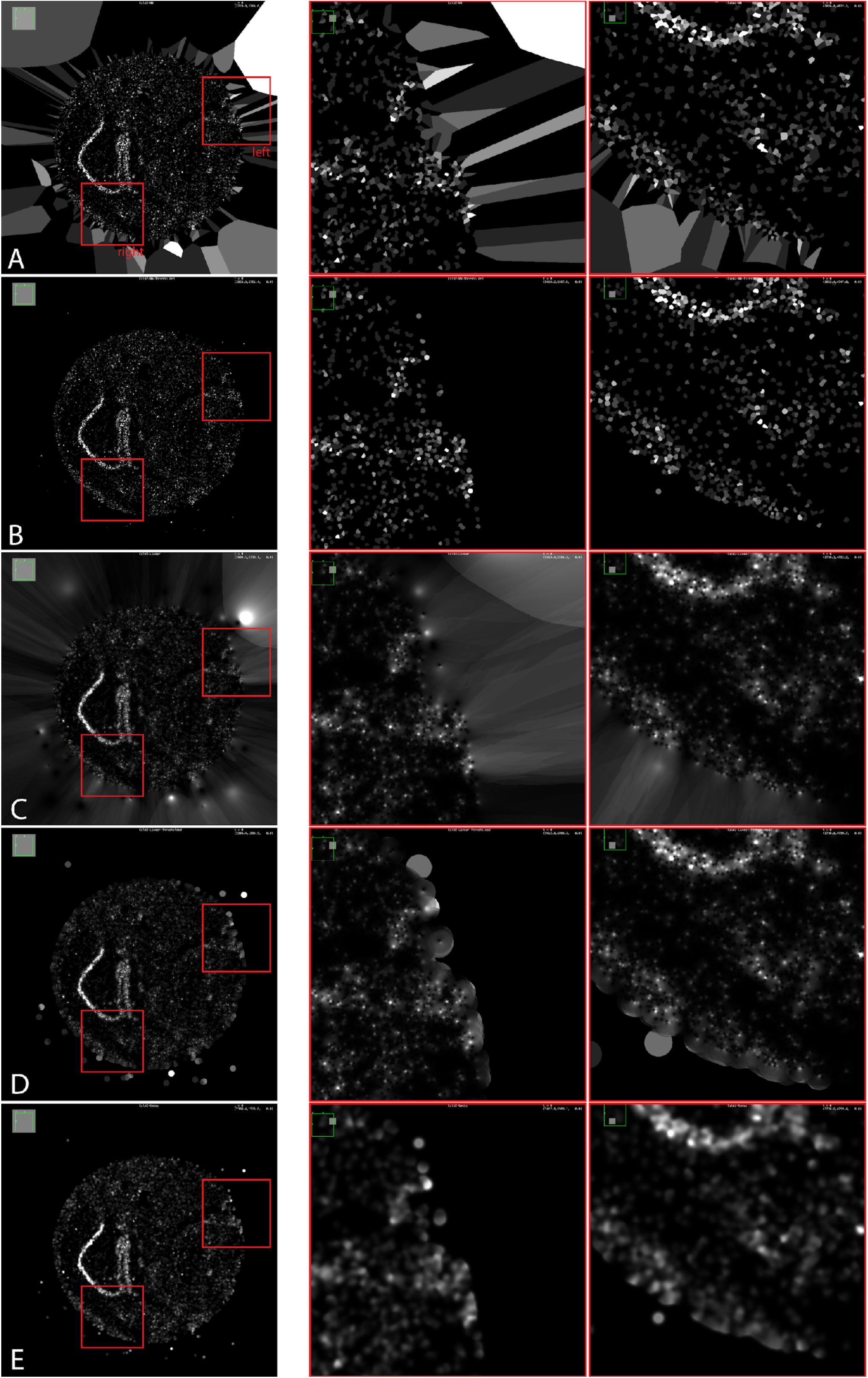
Comparison of different point cloud rendering methods supported by STIM. Each row shows a different rendering method for the same field of view of the gene *Calm2* on the left side, and two zoom-ins on the right side. (**A**) illustrates nearest-neighbor rendering. (**B**) show nearest neighbor rendering with a cut-off after the median distance between all sequenced locations (∼16 distance units). (**C**) shows distance-weighted rendering using a maximum number of 20 neighbors and quadratic distance weight. (**D**) shows the same distance-weighted rendering, but with an additional cut-off at 5x the median distance between all sequenced locations. (E) shows the Gaussian-distance weighted rendering with a sigma equivalent to the median distance between all sequenced locations. This rendering method was used for all experiments, figures and videos in this publication. The code for creating these representations can be found in the `net.imglib2` package.

**Supplementary Figure 7:**
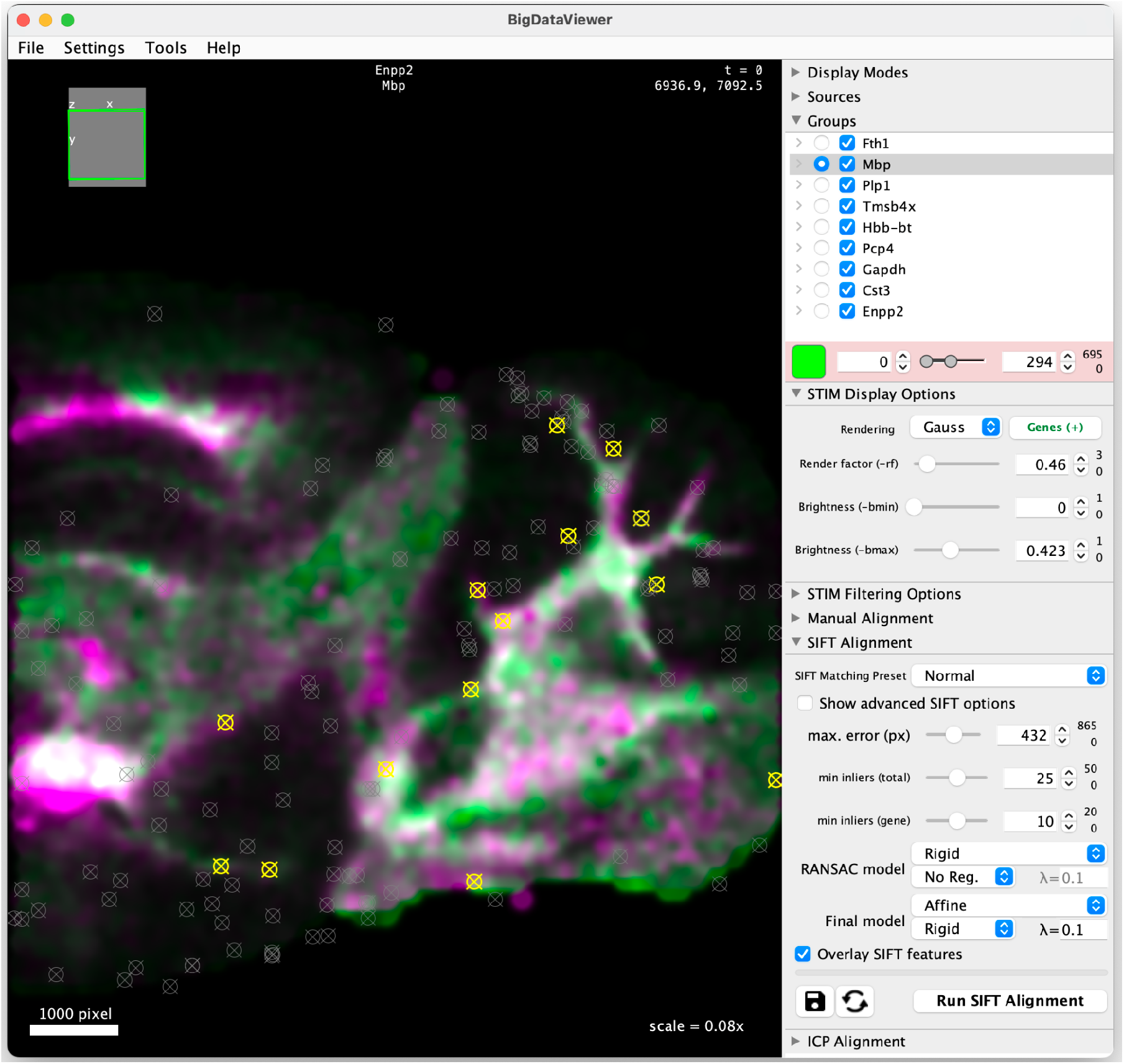
Interactive alignment using SIFT in the STIM BigDataViewer-based GUI. Corresponding SIFT features were identified using a rigid model and the final transformation computed from the features was a rigidly regularized affine model. Yellow crosshairs indicate automatically identified corresponding features in the visible gene (*Mbp*), while gray crosshairs show corresponding features in the other genes that were also used for alignment (*Fth1, Plp1*-*Enpp2*). The GUI allows to interactively modify visualization parameters, filtering, manual alignment, SIFT alignment and ICP alignment (Note: ICP iterations are interactively updated).

**Supplementary Table 1:** Comparison of 3D registration methods for Spatial Transcriptomics data. We selected properties that are important for aligning the Open-ST metastatic lymph node, a large dataset containing ∼1 million cells (19 sections spanning ∼3,000×4,000×350 μm^3^, ∼60,000 cells/section). For methods using specific genes for alignment, we selected the top 10 highly variable genes (scanpy, using flavor ‘seurat’ with default parameters). *Visualization*: seamless interactive exploration of the data in 3D, i.e., via custom visualization tools or functions connecting to existing ones. *✓✓*: visualization and interactive alignment are available. *✓*: only visualization tools are available. *✗*: visualization tools not available. *Global*: the model’s ability to perform global registration of stacks, to regularize the propagation of errors after pairwise alignment across the final z-stack. *Non-rigid:* whether the method provides non-rigid transformation models. *CPU Time*: excluding time required for converting the dataset, manual selection of points, or tasks not strictly related to alignment. *Memory:* peak RAM used during alignment, excluding preprocessing tasks. *Automated*: the method does not require manual selection of *landmarks*, or extensive preprocessing and annotation of cell types or *regions*. Morpho: from *spateo*; Sc3D: using the sc3d mode (instead of wrapper for PASTE). For the methods based on region selection, we used the transcriptomic cluster identities from the original publication. Under *CPU time* and *Memory*, the best method is **bold**, and the second best is <u>underlined</u>. The following software versions were used: STIM (0.3.0) PASTE2 (1.0.1), spateo-release (1.1.0), GPSA (0.8), SPACEL (1.1.7), ST-GEARS (1.0.0). The benchmark was run on a quad-socket server equipped with 4x Intel(R) Xeon(R) Platinum 8280 CPUs and 4,227 GB of RAM. All tools were run using a single thread, to discard additional sources of overhead. Jobs requiring more than 1 day to run were terminated, thus did not finish (DNF).

| Method | Visualization | Global | Non-Rigid | CPU time | Memory | Automated |
| --- | --- | --- | --- | --- | --- | --- |
| STIM | ✓✓ | ✓ | ✗ | ~181 min | ~6.8 GB | ✓ |
| Morpho <sup>1</sup> | ✓ | ✗ | ✓ | >1 day (DNF) | ≥416 GB | ✓ |
| GPSA <sup>2</sup> | ✗ | ✗ | ✓ | >1 day (DNF) | ≥162 GB | ✓ |
| PASTE2 <sup>3</sup> | ✗ | ✗ | ✓ | >1 day (DNF) | ≥231 GB | ✓ |
| SPACEL <sup>4</sup> | ✗ | ✗ | ✓ | ~433 min | <u>~18 GB</u> | ✗ (clusters) |
| ST-GEARS <sub>5</sub> | ✓ | ✗ | ✓ | >1 day (DNF) | ≥358 GB | ✗ (clusters) |
| Sc3d <sup>6</sup> | ✓ (napari) | ✗ | ✗ | <u>~191 min</u> | ~24 GB | ✗ (clusters) |
| Semla <sup>7</sup> | ✓ | ✗ | ✗ | N/A | N/A | ✗ (landmarks) |
| STAlign <sup>8</sup> | ✗ | ✗ | ✓ | N/A | N/A | ✗ (landmarks) |
| Eggplant <sup>9</sup> | ✗ | ✗ | ✓ | N/A | N/A | ✗ (landmarks) |

#### Workflow for the alignment of the 10x Visium dataset

~~~
./st-resave
       -i /home/10x-Visium/section1_locations.csv,/home/10x-Visium/section1_reads.csv,sec1
       -i /home/10x-Visium/section2_locations.csv,/home/10x-Visium/section2_reads.csv,sec2
       -c /home/10x-Visium.n5
./st-explorer -i /home/10x-Visium.n5/
./st-align-pairs -c /home/10x-Visium.n5/ -n 100 -rf 0.5 --maxEpsilon 100 --minNumInliersGene 30
./st-align-global -c /home/10x-Visium.n5/ --absoluteThreshold 100 -rf 0.5 --lambda 0.0 --skipICP
./st-render -i /home/10x-Visium.n5/ -g Calm2 -rf 0.6 -s 0.1
./st-render -i /home/10x-Visium.n5/ -g Calm2 -rf 0.6 -s 0.1 –ignoreTransforms
~~~

#### 3D Rendering of the Slide-Seq dataset

In order to create the 3D rendering shown in **Fig. 2** (which is also **Supplementary Video 1**), we exported three genes of the aligned SlideSeq dataset in low (1609 × 1771 × 256 px) and high resolution (3217 × 3540 × 511 px) and merged both into an RGB image (*Calm2* is shown in white, *Ptdgs* in green, and *Mbp* in red).

We then created a 3D projection using the Fiji command “ `Image > Stacks > 3D project`” for both images, which creates a visually pleasing animation of both images.

In order to create the zoom-in effect we wrote a script that smoothly interpolates between the two datasets as they rotate, which can be found in the `examples.MakeMovie` class in the STIM repository (https://github.com/PreibischLab/STIM/blob/master/src/main/java/examples/MakeMovie.java).

## Notes

https://github.com/PreibischLab/STIM

https://github.com/rajewsky-lab/stimwrap

### Competing Interest Statement

This work is part of a larger patent application in which the authors are among the inventors. The patent application was submitted through the Technology Transfer Office of the Max-Delbrueck Center (MDC), with the MDC being the patent applicant.

### Summary of Updates

consolidated the manuscript, clear focus on the method; describe added features (interactive alignment of ST-data) and add benchmarks

